# Common predators and factors influencing their abundance in *Anopheles funestus* aquatic habitats in rural south-eastern Tanzania

**DOI:** 10.1101/2022.12.03.518970

**Authors:** Herieth H. Mahenge, Letus L. Muyaga, Joel D. Nkya, Khamis S. Kifungo, Najat F. Kahamba, Halfan S. Ngowo, Emmanuel W. Kaindoa

## Abstract

**Background:** The role that larval predators play in regulating the population of malaria vectors remains relatively unknown. This study aimed to investigate the common predators that co-exist with *Anopheles funestus* group larvae and evaluate factors that influence their abundance in rural south-eastern Tanzania.

**Methods:** Mosquito larvae and predators were sampled concurrently using standard dipper (350 ml) or 10 L bucket in nine villages in southern Tanzania. Predators were identified using standard identification keys. All positive habitats were geo-located and their physical features characterized. Water physicochemical parameters such as dissolved oxygen (DO), pH, electrical conductivity (EC), total dissolved solids (TDS) and temperature were also recorded.

**Results:** A total of 85 previously identified *An. funestus* aquatic habitats were sampled for larvae and potential predators. A total of 8,295 predators were sampled. Of these Coenagrionidae 57.7% (n=4785), Corixidae 12.8% (n=1,060), Notonectidae 9.9% (n=822), Aeshnidae 4.9% (n=405), Amphibian 4.5% (n=370), Dytiscidae 3.8% (n=313) were common. A total of 5,260 mosquito larvae were sampled, whereby *Anopheles funestus* group were 60.3% (n= 3,170), *Culex* spp. 24.3% (n= 1,279), *An. gambie s.l*. 8.3% (n= 438) and other anophelines 7.1% (n= 373). Permanent and aquatic habitats larger than 100m^2^ were positively associated with *An. funestus* group larvae (P<0.05) and predator abundance (P<0.05). Habitats with submerged vegetation were negative associated with *An. funestus* group larvae (P<0.05). Only dissolved oxygen (DO) was positively and significantly affect the abundance of *An. funestus* group larvae (P<0.05). While predators abundance were not impacted by all physicochemical parameters.

**Conclusion:** Six potential predator families were common in aquatic habitats of *An. funestus* larvae group. Additional studies are needed to demonstrate the efficacy of different predators on larval density and adult fitness traits. Interventions leveraging the interaction between mosquitoes and predators can be established to disrupt the transmission potential and survival of the *An. funestus* mosquitoes.

## Background

The control of larvae of malaria vectors forms part of integrated control [1]. Such interventions are effective, inexpensive and safe to non-target organisms [2–4]. They are, however, underused. A review conducted between 1987 and 2019 discussing the use of bacterial larvicides Derua *et al*., [5], were only able to describe 32 field based studies of these techniques. They suggested that larviciding should become more important as a vector control tool. However, a greater understanding of the larval biology is essential for effective application. In particular, the role played by predators in regulating the population of mosquito larvae remains relatively unstudied.

Many aquatic invertebrate predator species, including Aeshnidae [6], Notonectidae [6] and Dytiscidae [7] coexist with mosquito larvae [8]. Isolating predators and distinguishing them from other organisms can be done by direct observation of their behaviour, visual examination of their midgut contents, molecular methods or electrophoretic methods [9–11]. A single aquatic habitat may contain several species of predators. Such predators have been shown to be effective biocontrol agents against mosquitoes in different larval habitats [6,8,12]. For example, the mosquito larval mortality attributed to predators ranges between 54% and 90% depending on the environment, predator species diversity and density [8]. Although, mosquito predators directly, or indirectly, influence mosquito population dynamics [13], their effects on *An. funestus* dynamics are not understood. The use of such predators may limit mosquito larval abundance and reduce adult densities [8,14–16].

Although malaria is transmitted by several *Anopheles* species in Tanzania, *Anopheles funestus* is the primary vector [17–19]. This species is also highly resistant to common insecticides currently used for malaria control, in particular pyrethroids used in insecticide treated nets (ITNs) [20]. Despite a number of studies on the bionomics of these mosquitoes [21–23], and their aquatic habitats [24], the relationship between aquatic predators on *An. funestus* larval population remains unknown.

This study investigated the common aquatic predators and factors influencing their abundance in *An. funestus* aquatic habitats in south-eastern Tanzania. Specifically we aimed to 1) identify types of common predators co-existing with *An. funestus* group larvae in a rural part of Tanzania, 2) determine factors which might contributed to the abundance of these predators and 3) quantify the associations between different predator types and *An. funestus* group larval abundance.

## Methods

### Study area

A cross-sectional survey was conducted, between March and May 2022, in nine villages in south-eastern Tanzania, namely Chikuti (−8.6028°, 36.7288°), Mzelezi (−8.8934°, 36.7343°), Chirombola (−8.93041°, 36.75753°), Ebuyu (−8.9719°, 36.7608°), Mwaya (−8.91022°, 36.823139°) and Tulizamoyo (−8.35447°, 36.70546°) in Ulanga district and Ikwambi (−7.97927°, 36.81630°), Kisawasawa (−7.89657°, 36.88058°) and Mpofu (−8.17220°, 36.21651°) in Kilombero district (Figure 1). In this area *An. funestus* is responsible for more than 85% of overall malaria transmission [17]. The residents in these villages practise extensive rice farming, which creates suitable habitat for mosquito breeding. Common aquatic habitats for *An. funestus* in the villages are well known and have been previously characterized [24]. Eighty-five known habitats from the nine villages were sampled for both mosquito larvae and potential predators.

**Figure 1:**
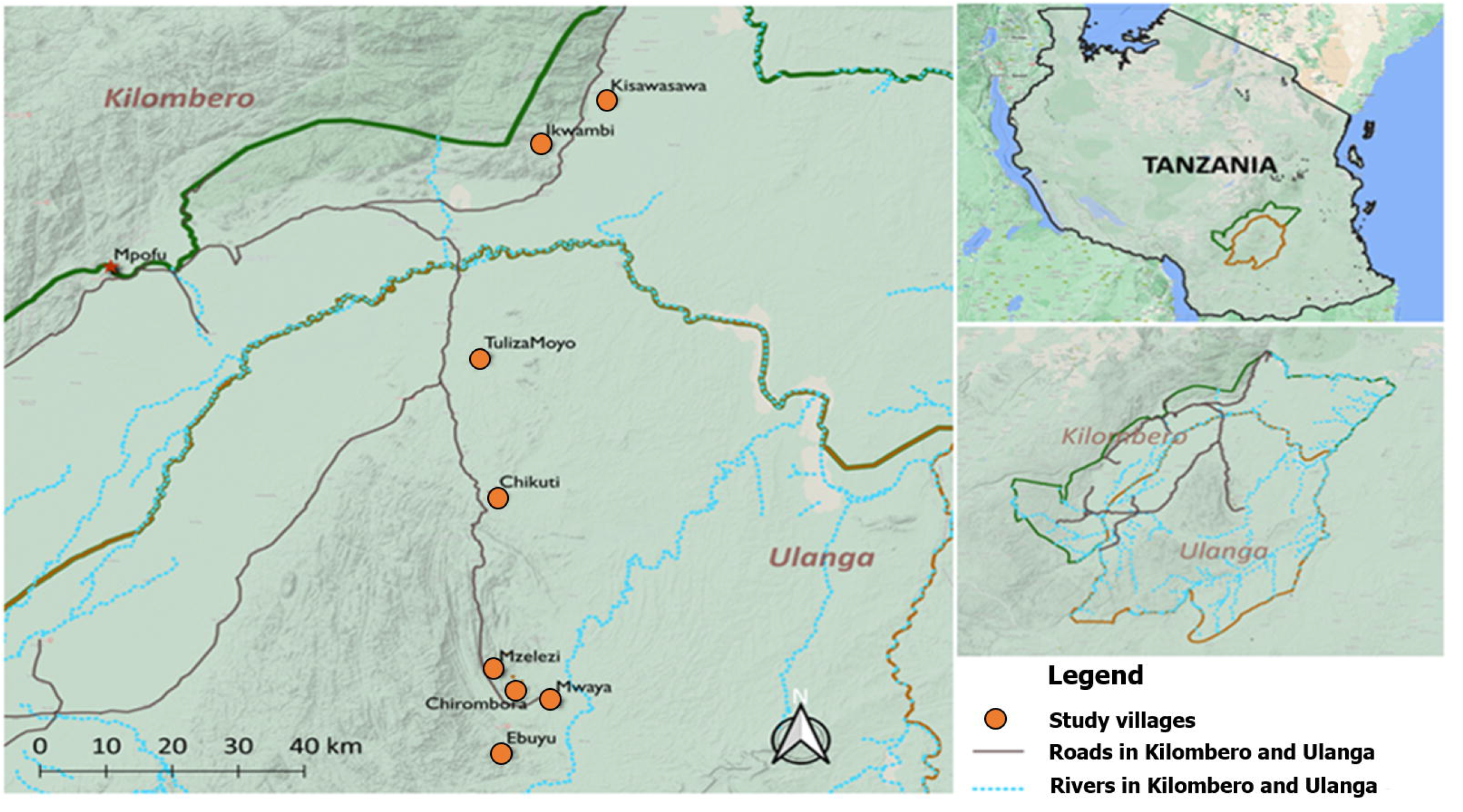
Map of Kilombero and Ulanga districts showing the nine study villages

**Figure 2:**
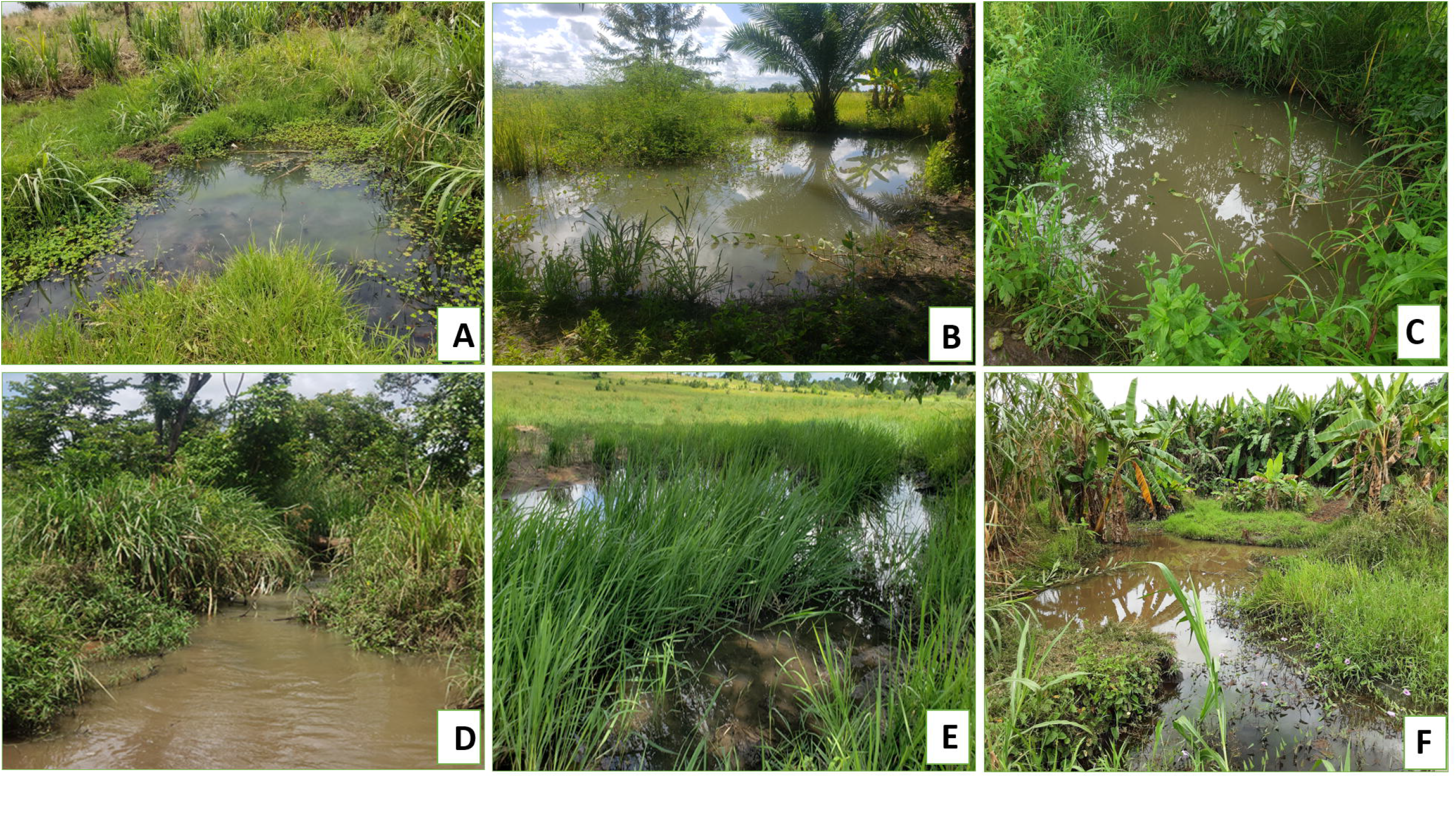
Different aquatic habitats which contain *Anopheles funestus* and predators; (**A**: Grounded pool, **B**: Brick/ sand pit, **C**: Man made well, **D**: River stream, **E**: Swamp, **F**: Grounded fed pool.

### Sampling and morphological identification of mosquito larvae and aquatic predators

Mosquito larvae and predators were sampled using standard dippers (350 ml) or 10 L buckets, as previously described [8,15,24]. A minimum of 3 dips and a maximum of 20 dips were taken depending on the size and depth of the habitat.

In a previous study mosquito larvae from the same villages were taken to the laboratory in Ifakara, allowed to emerge and eventually identified to species by PCR. Of those identified 53% were *An. funestus* s.s. whilst 28% were *An. rivulorum* and 12% were *An. leesoni* [24]. All three species were found to occupy the same habitats. A similar approach was followed with samples of fourth instar larvae during the present study but identification to species level was not performed. Earlier stage larvae were identified based on their predominant characteristics and separated into *An. funestus* group, *An. gambiae* s.l. or *Anopheles* sp. Culicines were identified to genera only. Predators were morphologically identified to family level using the keys of the Stroud Water Research Center [25] and Gerber and Gabriel [26]. Mosquito larvae and predators that were sampled by each dipper or bucket were counted and recorded. Additionally, geographical locations of the surveyed habitats were recorded at access points using a hand-held GPS device (Garmin eTrex 20x Handheld GPS Receiver).

### Aquatic habitats characterization

Although many individual dips were negative, all aquatic habitats sampled contained mosquito larvae. Their overall physical characteristics were recorded and physicochemical parameters of the water (pH, temperature, electrical conductivity (EC), total dissolved solids (TDS) were measured using a portable water quality meter (ZJ practical 4 in 1 Water Tester). A Trans Instruments Dissolved Oxygen Meter (HD3030) was used to measure dissolved oxygen (DO), using standard recording procedures. Habitats were classified as being either: swamp, stream, river, rice-field, stream-pool, ground-pool, ditch, spring-fed pool, puddle, hoof-print, man-made wells, brick or sand pit. Water colour was categorized as being clear (transparent and odourless) or coloured (cloudy, not transparent, turbid or with a film of oil).

The source of water was also classified as rainwater or others (non-rainwater). Algal quantities in the habitats were classified as none, moderate, or abundant. Algal type was classified as filamentous, green, blue-green or brown. Water was also classified as being stagnant, slow or fast moving. The land use surrounding the aquatic habitats was classified as scrub, cattle grazing or cultivated field. Shade over the habitats was classified as none, partial or heavy. Habitat size was measured using tape and classified as being less than 100 m^2^ or more than 100 m^2^. Vegetation quantity and vegetation type were also classified as (none, moderate or abundant) and (emergent, or submerged) respectively. Water bodies known to have existed for three months or more were considered to be permanent whilst other collections of water were considered to be ‘temporary’. Water depth was classified as being less than 50 cm or more than 50 cm deep. The distance from aquatic habitats to the nearest houses were estimated visually and classified as being less than 100 m or more than 100 m.

### Statistical analysis

Analysis was done using open source software R version 4.2.1. [27]. Generalised linear mixed effects models (GLMM) using template model builder (TMB) with zero-inflated negative binomial implemented under the *glmmTMB* package [28] were used to (i) assess the associations between water physicochemical parameters and the abundance of aquatic predators ii) assess the associations between water physicochemical parameters and the abundance of *An. funestus* group larvae (iii) assess which habitat characteristics contributed to the abundance of predators and *An. funestus* group larvae and (iv) assess the impact of each predator family on the abundance of *An. funestus* group larvae. All variables (i-iv) were assessed individually and later combined in the final model.

Due to a large number of dips with zero larvae the negative binomial with zero inflated models were used. In all models, the study villages in which the aquatic habitats were identified and habitat ID were used as random terms to capture unexplained variations between villages and habitats. The best fitting models were selected using Akaike Information Criterion (AIC) and results presented as risk ratios (RR) at 95% CI and statistical significance was considered when the P-value < 0.05.

## Results

### Distribution of *Anopheles funestus* larvae, other mosquito larvae and aquatic predators

A total of 85 aquatic habitats that contained *Anopheles funestus* group larvae were identified and characterized. In these habitats 8,295 predators belongs in were sampled. Among all sampled predators, only 7,906 predators were identified belonging to eight families. Of these, Coenagrionidae accounted for 57.7% (n=4785), Corixidae 12.8% (n=1,060), Notonectidae 9.9% (n=822), Aeshnidae 4.9% (n=405), Amphibian 4.5% (n=370), Dytiscidae 3.8% (n=313), Belostomatidae 1.2% (n=103) and Nepidae 0.6 % (n=48). Three hundred and eighty nine (4.6 % of the total invertebrates) were not identified due to lack of an appropriate key. A total of 5,260 larvae were collected, with *An. funestus* group larvae accounting for 60.3% (n=3,170) of the total, *Culex* spp 24.3 (n=1,279), *An. gambiae s.l*. 8.3% (n= 438), and other anopheline larvae 7.1% (n= 373). The predators from six family were more abundant and common in all aquatic habitats namely Coenagrionidae, Corixidae, Notonectidae, Aeshnidae, Amphibians and Dytiscidae (Table 2). The distribution of predators, *An. funestus* group larvae and other mosquito larvae per each habitats type is as presented in Table 2 and 3.

**Table 1:** Mean number and standard error (se) of different mosquito larvae species sampled from different aquatic habitats in Ulanga and Kilombero in Southern Eastern Tanzania

| Study site | Habitat information | Mean number and standard error (se) of different mosquitoes larvae |  |  |  |  |  |
| --- | --- | --- | --- | --- | --- | --- | --- |
| Habitat type |  | Total habitats | *An. funestus*<br>group | *An. gambiae s.l* | Other<br>anophelines | *Culex* spp | Total |
|  |  |  | Mean(2se) | Mean(2se) | Mean(2se) | Mean(2se) |  |
| Ulanga and Kilombero | Brick or sand pit | 12 | 37.8(19.64) | 2.7(2.5) | 3.2(6.0) | 15.8(11.9) | 713 |
|  | Ditch | 8 | 16.5(8.4) | 1.5(1.1) | 5.0(7.4) | 20.2(22.3) | 346 |
|  | Grounded pool | 1 | 19.0(NA) | 3.0(NA) | 0 | 12.0(NA) | 36 |
|  | Man- made wells | 20 | 11.2(3.4) | 3.8(5.3) | 0.6(0.6) | 5.6(3. 6) | 422 |
|  | Rice field | 2 | 40.5(51.9) | 3.0(3.9) | 10.5(20.6) | 24.5(34.3) | 157 |
|  | River stream | 34 | 61.7(29.8) | 9.0(8.0) | 5.4(4.7) | 13.3(10.2) | 3039 |
|  | Spring- fed pool | 2 | 25.0(17.6) | 0 | 0 | 29.5(38.2) | 109 |
|  | Swamp | 6 | 18.8(4.6) | 0.7(1.0) | 13.2(10.8) | 40.7(19.4) | 440 |
|  | **Total** | 85 | 3170 | 438 | 373 | 1279 | 5260 |

**Table 2:** Mean number and standard error (se) of different predators sampled from different aquatic habitats in Ulanga and Kilombero in Southern Eastern Tanzania

| Study site | Habitat information |  | Mean number and standard error of different predators |  |  |  |  |  |  |  |  |
| --- | --- | --- | --- | --- | --- | --- | --- | --- | --- | --- | --- |
| Ulanga and Kilombero | Habitat type | Total habitats (N) | Aeshnidae | Coenagrionidae | Dytiscidae | Notonectidae | Corixidae | Nepidae | Belomastidae | Amphibians | Unidentified species |
|  |  |  | Mean(2se) | Mean(2se) | Mean(2se) | Mean(2se) | Mean(2se) | Mean(2se) | Mean(2se) | Mean(2se) | Mean(2se) |
|  | Brick or sand pit | 12 | 8.25(6.07) | 33.0(19.42) | 3.83(2.61) | 10.33(6.0) | 19.42(22.95) | 0.25(0.35) | 1.42(1.59) | 2.08(1.35) | 2.0(2.84) |
|  | Ditch | 8 | 1.0(1.11) | 10.3(6.7) | 7.63(7.27) | 6.75(9.3) | 1.13(0.94) | 0.25(0.49) | 3.25(5.58) | 23.5(45.78) | 2.25(1.92) |
|  | Grounded pool | 1 | 0.0(0.0) | 46.0(NA) | 0.0(0.0) | 0.0(0.0) | 4.0(0.0) | 0.0(0.0) | 0.0(0.0) | 1(NA) | 0.0(0.0) |
|  | Man-made wells | 20 | 7.05(6.38) | 32.2(33.35) | 2.45(2.53) | 23.75(18.18) | 5.95(6.89) | 0.55(0.78) | 0.05(0.10) | 1.45(1.49) | 0.90(0.98) |
|  | Rice field | 2 | 2.50(2.94) | 37.5(42.14) | 9.0(17.64) | 5.0(9.8) | 1.0(9.8) | 1.00(1.96) | 0.0(0.0) | 1.00(0.00) | 16.0(31.36) |
|  | River stream | 34 | 1.50(0.98) | 97.18(24.1) | 2.0(1.5) | 1.21(0.89) | 17.4(11.0) | 0.76(0.63) | 1.65(1.33) | 3.44(2.94) | 6.76(4.06) |
|  | Spring-fed pool | 2 | 17.5(22.54) | 46.5(46.07) | 0.5(1.0) | 36.5(2.94) | 2.0(1.96) | 1.00(0.0) | 0.0(0.0) | 2.0(1.96) | 1.0(NA) |
|  | Swamp | 6 | 11.0(7.62) | 24.17(13.71) | 11.7(12.0) | 7.5(7.4) | 16.17(15.48) | 0.33(0.41) | 0.50(0.98) | 0.67(0.65) | 10.83(4.54) |
| **Ulanga** | **Total** | 85 | 405 | 4785 | 313 | 822 | 1060 | 48 | 103 | 370 | 389 |

**Table 3:** Number of each aquatic habitats type showing the co-existence of different mosquito larvae group

| Study site | Habitat information |  | Number of habitats with different mosquitoes species |  |  |  |
| --- | --- | --- | --- | --- | --- | --- |
|  | Habitat type | Total habitats | Habitats with *An. funestus s.l* | Habitats with *An. gambiae s.l* | Habitats with other anopheline | Habitats with *Culex spp* |
| Ulanga and Kilombero | Brick or sand pit | 12 | 12 | 7 | 2 | 10 |
|  | Ditch | 8 | 8 | 5 | 2 | 5 |
|  | Grounded pool | 1 | 1 | 1 | 0 | 1 |
|  | Man-made wells | 20 | 20 | 10 | 4 | 13 |
|  | Rice field | 2 | 2 | 2 | 1 | 2 |
|  | River stream | 34 | 33 | 16 | 9 | 21 |
|  | Spring- fed pool | 2 | 2 | 0 | 0 | 2 |
|  | Swamp | 6 | 6 | 2 | 5 | 6 |
|  | **Total** | **85** | **85** | **46** | **23** | **60** |

### Water physicochemical parameters and their influence on the abundance of predators and *An. funestus* larvae group

There was no apparent association between physicochemical parameters (pH, temperature, TDS, EC and DO) and predator abundance (P>0.05, Table 9). Temperature, pH, TDS, EC also had no impact on the abundance of *An. funestus* group larvae but DO was positively associated with the abundance of *An. funestus* group larvae (P<0.05, Table 8).

### Characteristics of aquatic habitats and their influence on the abundance of *An. Funestus* larval group and predators

*Anopheles funestus* group larvae and predators were found in high abundance in habitats larger than 100 m^2^ and at the edges of streams and rivers (habitats with fast moving water) (P<0.05, tables 4 and 5), whilst low abundance of larvae was associated with habitats with submerged vegetation (P<0.05, Table 4). Other aquatic habitat characteristics including algae quantity and type, shade over the habitats, water depth, vegetation quantity, environment surrounding the aquatic habitats and the distance from the nearest houses were found to have no impact on *An. funestus* group larval abundance and predator abundance.

**Table 4:**
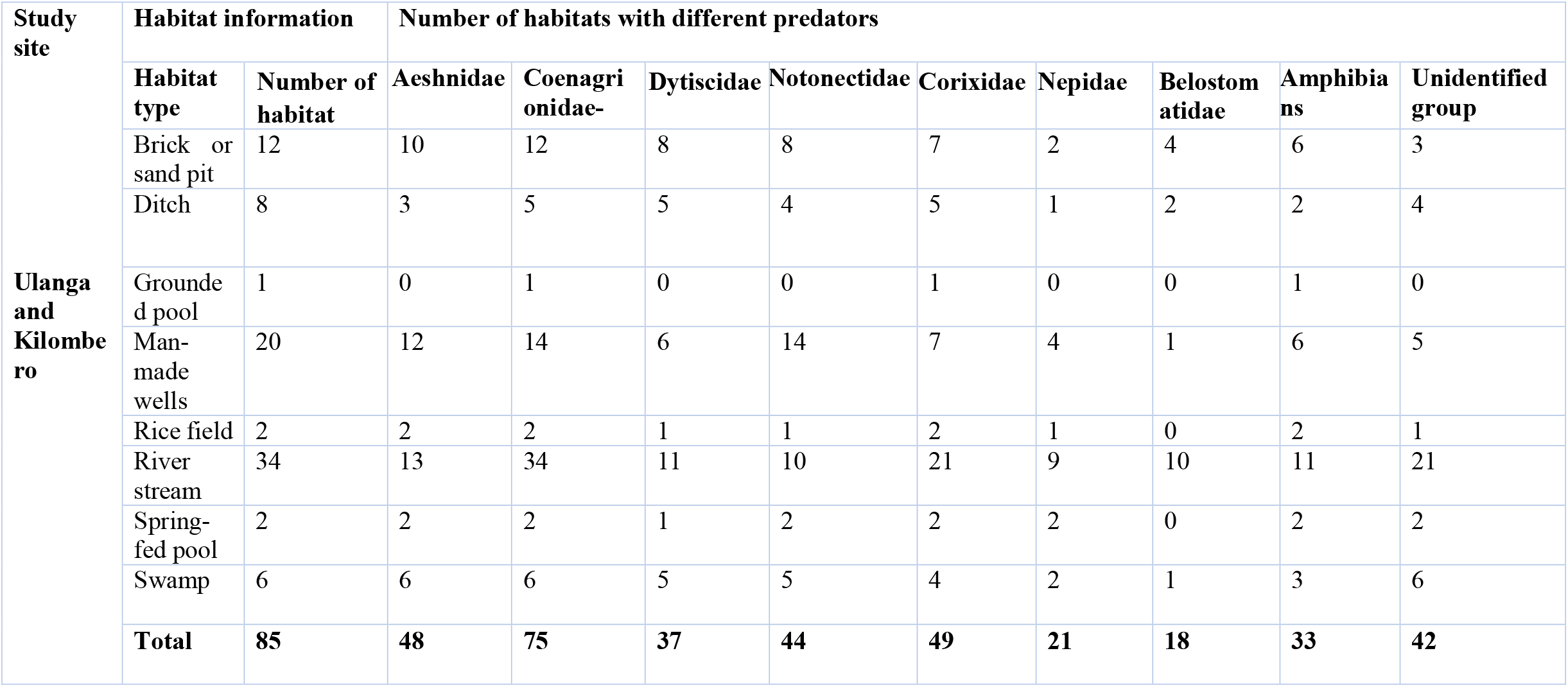
Number of each aquatic habitats type showing the co-existence of different predators

| Study site | Habitat information |  | Number of habitats with different predators |  |  |  |  |  |  |  |  |
| --- | --- | --- | --- | --- | --- | --- | --- | --- | --- | --- | --- |
|  | Habitat type | Number of habitat | Aeshnidae | Coenagri<br>onidae- | Dytiscidae | Notonectidae | Corixidae | Nepidae | Belostom<br>atidae | Amphibia<br>ns | Unidentified<br>group |
| Ulanga<br>and<br>Kilombe<br>ro | Brick or<br>sand pit | 12 | 10 | 12 | 8 | 8 | 7 | 2 | 4 | 6 | 3 |
|  | Ditch | 8 | 3 | 5 | 5 | 4 | 5 | 1 | 2 | 2 | 4 |
|  | Grounded pool | 1 | 0 | 1 | 0 | 0 | 1 | 0 | 0 | 1 | 0 |
|  | Man-made<br>wells | 20 | 12 | 14 | 6 | 14 | 7 | 4 | 1 | 6 | 5 |
|  | Rice field | 2 | 2 | 2 | 1 | 1 | 2 | 1 | 0 | 2 | 1 |
|  | River<br>stream | 34 | 13 | 34 | 11 | 10 | 21 | 9 | 10 | 11 | 21 |
|  | Spring-fed pool | 2 | 2 | 2 | 1 | 2 | 2 | 2 | 0 | 2 | 2 |
|  | Swamp | 6 | 6 | 6 | 5 | 5 | 4 | 2 | 1 | 3 | 6 |
|  | **Total** | **85** | **48** | **75** | **37** | **44** | **49** | **21** | **18** | **33** | **42** |

**Table 5:** Univariate and multivariate regression analysis of different aquatic habitat characteristics and their association with the abundance of *Anopheles funestus* larvae

| Aquatic habitat | Univariate analysis |  | Multivariate analysis |  |
| --- | --- | --- | --- | --- |
|  | RR (95% LC, UC) | P-values | RR (95% LC, UC) | P-values |
| **Algae quantity** |  |  |  |  |
| None | 1 |  | 1 |  |
| Moderate | 1.21 [0.66, 2.20] | 0.535 | 1.44 [0.76, 2.73] | 0.260 |
| Abundant | 1.94 [0.89, 4.20] | 0.094 | 1.72 [0.72, 4.11] | 0.224 |
| **Habitat size** |  |  |  |  |
| Less than 100 M | 1 |  | 1 |  |
| Greater than 100M | 2.54 [1.65, 3.89] | < 0.05 | 2.56 [1.58, 4.14] | < 0.05 |
| **Vegetation type** |  |  |  |  |
| None | 1 |  |  |  |
| Emergent | 0.73 [0.32, 1.63] | 0.435 | 0.64 [0.35, 1.18] | 0.151 |
| Sub-merged | 0.21 [0.04, 1.08] | 0.062 | 0.10 [0.02, 0.57] | < 0.05 |
| **Water Movement** |  |  |  |  |
| Stagnant | 1 |  | 1 |  |
| Slow | 1.45 [0.90, 2.35] | 0.127 | 1.29 [0.76, 2.18] | 0.343 |
| Fast | 1.82 [0.96, 3.45] | 0.064 | 2.79 [1.39, 5.63] | < 0.05 |
| **Shade over habitat** |  |  |  |  |
| None | 1 |  | 1 |  |
| Partial | 0.94 [0.55, 1.61] | 0.818 | 1.13 [0.64, 2.00] | 0.675 |
| **Water depth** |  |  |  |  |
| Less than 50 cm | 1 |  | 1 |  |
| More than 50 cm | 1.59 [1.07, 2.38] | 0.02 | 0.94 [0.56, 1.59] | 0.824 |
| **Water type** |  |  |  |  |
| Temporary | 1 |  | 1 |  |
| Permanent | 1.18 [0.76, 1.82] | 0.456 | 0.77 [0.44, 1.33] | 0.343 |
| **Environment around habitat** |  |  |  |  |
| Cultivated field | 1 |  | 1 |  |
| Scrub | 1.48 [0.94, 2.31] | 0.089 | 1.30 [0.80, 2.11] | 0.282 |
| **Water colour** |  |  |  |  |
| Turbid | 1 |  | 1 |  |
| Clear | 0.99 [0.58, 1.71] | 0.975 | 0.77 [0.46, 1.31] | 0.338 |
| **Distance from houses** |  |  |  |  |
| Less than 100 | 1 |  |  |  |
| More than 100 M | 1.28 [0.79, 2.06] | 0.320 | 1.02 [0.60, 1.75] | 0.937 |

### Association of different predators with the abundance of *An. funestus* group larvae

Coenagrionidae and Dytiscidae were positively associated with *An. funestus* group larval abundance (P<0.05, Table 10) whilst Notonectidae and Corixidae were negatively associated with *An. funestus* abundance (P<0.05, Table 10). No strong association between abundance of *An. funestus* group larval abundance and some predator families including Aeshnidae and Belostomatidae were found (P>0.05, Table 10).

## Discussion

Ecological interactions such as predation and competition are key drivers of population size of numerous organisms [29]. In the context of mosquito borne diseases, predators play an important role in regulating the diseases transmitting mosquitoes directly through feeding on mosquito larvae or indirectly through compromising mosquito fecundity, growth rate and growth trajectories [13,29]. Also, they regulate the *Anopheles* populations naturally through predation, parasitism and competition [30], the use of aquatic predators represents a potentially simple and practical biological technology for the control of disease transmitting mosquitoes [8]. Biological control methods, including the use of naturally occurring predators, have been utilised for vector control in many parts of the world [14,31,32].

The present study was undertaken to investigate the common predators and factors influencing their abundance in *An. funestus* aquatic habitats. Eight different families of predators co-existing with *An. funestus* group were identified, six of which, namely; Coenagrionidae, Corixidae, Notonectidae, Aeshnidae, Amphibians, and Dytiscidae were common in all habitat types (Table 2). Similarly, previous studies confirmed different predominant family Coenagrionidae [33], Dytiscidae [33], Notonectidae [34] in mosquitoes larval habitats. In addition, Notonectidae shown to have direct or direct effects on mosquito larvae population [35]. Current study found consistently, high mean number of Coenagrionidae family in all habitats type as compare to other predators families (Table 2). This implies that the characteristics of the surveyed aquatic habitats were favouring the survival and growth of these predators, hence led to high abundance.

Variation and abundance of different predators across different aquatic habitats were strongly associated with the some physical characteristics of the habitat. The high abundance of predators were generally observed in permanent habitats with fast moving water and larger than 100m^2^, e.g. brick or sand pits, man-made wells, river streams and swamps, similar to observations from other settings [16,36–40]. Such permanent aquatic habitats contain favourable amounts of both decomposed organic and inorganic matter which serve as food for predators, and these habitats allow colonization of the predators than temporal and simple structural habitats [41].

Interestingly, it was noted that aquatic habitats larger than 100m^2^ with fast moving water were positively associated with the abundance of *An. funestus* group larvae. Aquatic habitats with submerged vegetation were negatively associated with the abundance of *An. funestus* group larvae (Table 5). Previous studies have described a positive association between aquatic habitats with emergent vegetation and abundance of *An. funestus* group larvae [24,42], but not a negative association between abundance and habitats with submerged vegetation. This may be due to the season in which sampling was undertaken may have an influence on the nature of the vegetation in the aquatic habitats and movement of water.

With the exception of dissolved oxygen (DO), there was no association between other water physicochemical parameters and *An. funestus* larvae group abundance. On the other hand, predator abundance was not impacted by any of the measured water physicochemical parameters. As in a *previous study*, dissolved oxygen was found to be positively associated with the abundance of *An. funestus* group larvae [43]. This may be due to the preference of *An. funestus* larvae to breed in fresh and clear water which contains high levels of dissolved oxygen.

The current results are in line with the findings reported by *Bashar et al*., [44], which indicated 308 that dissolved oxygen is the preeminent predictor for the abundance of *Anopheles* mosquito 309 larvae in aquatic habitat Several factors, such as physical, chemical, biological and microbiological processes influence the levels of dissolved oxygen concentration in water, such that low dissolved oxygen concentrations, < 3 mg/L in fresh water indicate high level of pollution [33]. In this study the mean of dissolved oxygen was 6.2 mg/L and ranges from 1.12-16.56 mg/L (Table 7), this indicates that most of these aquatic habitats contained the highest amount of dissolved oxygen and aeration which favoured the abundance of the *An. funestus* group larvae and predators.

**Table 6:** Univariate and multivariate regression analysis of different aquatic habitat characteristics and their association with the abundance of predators

| Aquatic habitat | Univariate analysis |  | Multivariate analysis |  |
| --- | --- | --- | --- | --- |
|  | RR (95% LC, UC) | P-values | RR (95% LC, UC) | P-values |
| **Habitat size** |  |  |  |  |
| Less than 100 M | 1 |  | 1 |  |
| Greater than 100M | 2.92 [1.72, 4.96] | <0.05 | 3.52 [1.90, 6.53] | < 0.05 |
| **Vegetation type** |  |  |  |  |
| None | 1 |  |  |  |
| Emergent | 0.75 [0.37, 1.50] | 0.409 | 1.17 [0.53, 2.56] | 0.798 |
| Submerged | 0.42 [0.07, 2.54] | 0.346 | 0.43 [0.08, 2.27] | 0.323 |
| **Water Movement** |  |  |  |  |
| Stagnant | 1 |  | 1 |  |
| Slow | 2.09 [1.22, 3.58] | <0.05 | 0.98 [0.52, 1.87] | 0.961 |
| Fast | 2.91[1.39, 6.13] | <0.05 | 2.25 [0.94, 5.39] | < 0.05 |
| **Shade over habitat** |  |  |  |  |
| None | 1 |  | 1 |  |
| Partial | 1.25 [0.64, 2.43] | 0.508 | 1.66 [0.77, 3.56] | 0.192 |
| **Water depth** |  |  |  |  |
| Less than 50 cm | 1 |  | 1 |  |
| More than 50 cm | 2.03 [1.17, 3.51] | <0.05 | 1.12 [0.58, 2.18] | 0.727 |
| **Water type** |  |  |  |  |
| Temporary | 1 |  | 1 |  |
| Permanent | 2.64 [1.48, 4.69] | <0.05 | 2.89 [0.99, 4.41] | <0.05 |
| **Environment around habitat** |  |  |  |  |
| Cultivated field | 1 |  | 1 |  |
| scrub | 1.21 [0.71, 2.06] | 0.477 | 1.06 [0.57, 1.98] | 0.851 |
| **Water colour** |  |  |  |  |
| Coloured | 1 |  | 1 |  |
| Clear | 1.44 [0.56, 3.67] | 0.447 | 0.86 [0.44, 1.68] | 0.651 |
| **Distance from home** |  |  |  |  |
| Less than 100 M | 1 |  |  |  |
| More than 100 M | 1.164 [0.64, 2.12] | 0.620 | 0.72 [0.35, 1.46] | 0.364 |

**Table 7:** Mean and range values of water physicochemical parameters in the aquatic habitats

| Water characteristics | Mean ( Range) |
| --- | --- |
| pH | 6.3 [5.70-7.82] |
| Temperature (<sup>0</sup>C) | 28.0 [23.3-36.] |
| TDS (ppm) | 126.8 [23.0-395.0] |
| EC (μS/cm) | 253.7 [40.1-619.0] |
| DO (mg/L) | 6.2 [1.12-16.56] |

Water pH is one of the important factors for aquatic organisms [45]. It can limit the abundance and distribution of aquatic organisms because it is directly related to their cellular functions [45], growth and development as well as their survival [45,46]. It has been noted that mosquitoes can tolerate extremely high levels of water pH. The current study shows that pH was not apparently associated with the abundance of either *An. funestus* group larvae or predators in the aquatic habitats (Table 8 & 9). This correlates with Akeju *et al*., [47], Obi *et al* [48] and Chaiphongpachara *et al*., [33,49] and suggests that *An. funestus* group larvae and predators are ble to tolerate a wide range of pH in different environments. In addition, the current study shows the range of pH in the aquatic habitats was 5.70 to 7.82 (Table 7). These results are in line with the previous findings which shows the association of *Anopheles* larvae with aquatic insects including predators in a wide range of pH concentration in their aquatic habitats [33,50]. Both mosquito larvae and aquatic insects including predators have the328 mechanisms that enable them to inhabit such environments [45].

**Table 8:**
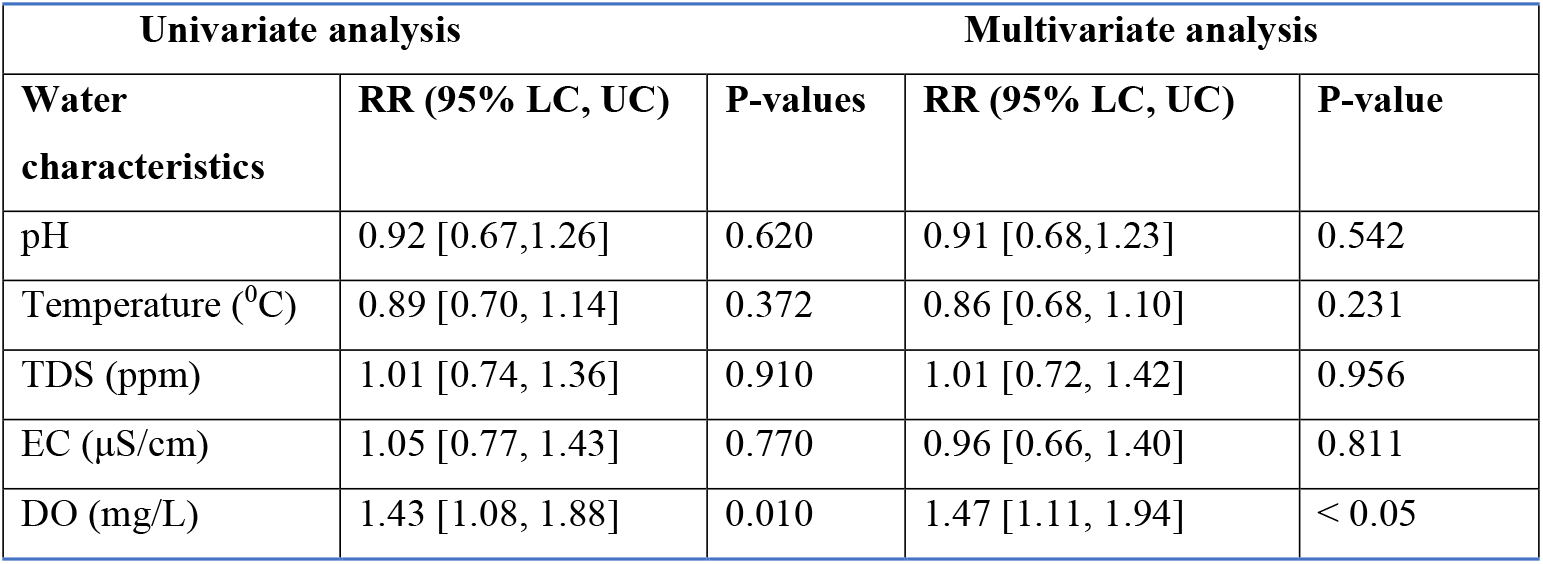
Univariate and multivariate analysis of associations between water physicochemical parameters and the abundance of *Anopheles funestus* larvae

| Univariate analysis |  |  | Multivariate analysis |  |
| --- | --- | --- | --- | --- |
| Water characteristics | RR (95% LC, UC) | P-values | RR (95% LC, UC) | P-value |
| pH | 0.92 [0.67,1.26] | 0.620 | 0.91 [0.68,1.23] | 0.542 |
| Temperature (<sup>0</sup>C) | 0.89 [0.70, 1.14] | 0.372 | 0.86 [0.68, 1.10] | 0.231 |
| TDS (ppm) | 1.01 [0.74, 1.36] | 0.910 | 1.01 [0.72, 1.42] | 0.956 |
| EC (μS/cm) | 1.05 [0.77, 1.43] | 0.770 | 0.96 [0.66, 1.40] | 0.811 |
| DO (mg/L) | 1.43 [1.08, 1.88] | 0.010 | 1.47 [1.11, 1.94] | < 0.05 |

**Table 9:** Univariate and multivariate analysis of associations between water physicochemical parameters and the abundance of predators in *Anopheles funestus* aquatic habitats

| Univariate analysis |  |  | Multivariate analysis |  |
| --- | --- | --- | --- | --- |
| Water characteristics | RR (95% LC, UC) | P-values | RR (95% LC, UC) | P-value |
| pH | 1.00 [0.74, 1.39] | 0.981 | 1.17 [0.84, 1.64] | 0.356 |
| Temperature (<sup>0</sup>C) | 0.85 [0.59, 1.23] | 0.390 | 0.84 [0.57, 1.24] | 0.381 |
| TDS (ppm) | 1.31 [0.99, 1.74] | 0.056 | 1.44 [0.98, 2.13] | 0.064 |
| EC (μS/cm) | 1.17 [0.86, 1.58] | 0.312 | 0.83 [0.53, 1.31] | 0.432 |
| DO (mg/L) | 1.25 [0.93, 1.68] | 0.142 | 1.31 [0.93, 1.85] | 1.118 |

**Table 10:** Univariate and multivariate regression analysis of different aquatic predators and their association with the abundance of *Anopheles funestus*

| Aquatic predators | Univariate analysis |  | Multivariate analysis |  |
| --- | --- | --- | --- | --- |
|  | RR (95% LC, UC) | P-values | RR (95% LC, UC) | P-value |
| **Aeshnidae** | 1.007 [0.984, 1.031] | 0.524 | 1.014 [0.989,1.039] | 0.268 |
| **Coenagrionidae** | 1.008 [1.005, 1.011] | <0.05 | 1.008 [1.006,1.011] | <0.05 |
| **Dytiscidae** | 1.043 [1.007, 1.078] | <0.05 | 1.035 [1.006,1.065] | <0.05 |
| **Notonectidae** | 0.998 [0.990, 1.006] | 0.685 | 0.988 [0.978,0.998] | <0.05 |
| **Corixidae** | 0.994 [0.984, 1.005] | 0.274 | 0.990 [0.984,0.997] | <0.05 |
| **Belomastidae** | 0.99 [0.921, 1.069] | 0.794 | 0.973 [0.920,1.028] | 0.327 |

Temperature is an important factor mediating predators and mosquito larvae interactions [51]. For example, it affects the ecology, physiology, metabolic processes and overall fitness of organisms [52]. Implication on the interaction between predators and mosquitoes as well as their behaviour performance in the aquatic habitats is mediated by temperature, because temperature plays an essential role as a regulatory mechanism that drives both physiological and biochemical activities [53]. Both *An. funestus* group larvae and predators share the same aquatic habitats which their temperature ranged from 23.3-36.4°C (Table 7), this shows that *An. funestus* and predators preferred warm conditions for their survival, development and colonization. The findings of the current study are in line with findings by Dida *et al*., which reported that both predators and prey preferred temperatures above 18°C and above 25°C [33].

Temperature was not significantly associated with the abundance of *An. funestus* group larvae and predators in the aquatic habitats which correlates previous findings [24], however some studies reported contrary findings showing positive association between *An. funestus* larvae and by temperature [43,44,47,54]. Most studies have mainly focused on the impacts of 18 terrestrial temperature on mosquitoes but a 344 limited number of studies focussed on aquatic habitats in the context of thermal tolerance, particularly for vector mosquitoes and their predators. This necessitates further investigations across seasons.

While studies done elsewhere yielded evidence that electrical conductivity is positively associated with the abundance of *An. funestus* larvae [47,54]. Further study found that higher levels of electrical conductivity was due to the application of agricultural fertilisers, pesticides and herbicides [55], but this study did not find any significant association between electrical conductivity and abundance of both, *An. funestus* larvae group and predators in the aquatic habitats. Electrical conductivity ranged between 40.1-619.0 μS/cm (Table 7),this shows *An. funestus* group larvae and predators can survive in a wide range of electrical conductivity in the aquatic habitats which similar to report by *Dida et al*, which suggested mosquito larvae and predators were most preferable in the aquatic habitats with electrical conductivity ranges between 162.9μS/cm and 166 μS/cm [33].

In aquatic habitats, higher total dissolved solids have harmful impacts on the aquatic organisms. It changes the mineral water contents, which is important for survival of predators and mosquitoes larvae. Furthermore, it determines the flow of water out of an organism’s cell. In this study, there was a wide range of total dissolved solids in the aquatic habitats in which it ranges between 23.0-395.0 ppm (Table 7). This variation might be the same as previously reported by another study that total dissolved solids in the aquatic habitat is highly dependent on the different factors such as the pattern use of different chemicals in the environments (like agriculture pesticides) [56]. Also, these results correlates to Oyewole *et al*., [57], but contrary to Abai *et al*., and Dida *et al*., which suggested that *Anopheles* mosquito associated with very high total dissolved solids (1,261.40 ± 1,214.31) [58] or very low 8–87 ppm [33].

This study revealed that various predator family share similar aquatic habitats, whereby Coenagrionidae were found in 75 habitats, Aeshnidae, in 48 habitats, Corixidae in 49 habitats, Notonectidae in 44 habitats, Dytiscidae in 37 habitats, Nepidae in 21 habitats, Amphibian in 33 habitats, Belostomatidae in 18 habitats and unidentified group in 42 habitats (Table 4). Furthermore, the results show that among 85 *An. funestus* habitats, a total of 46 were co-inhabited by *An. gambiae s.l*, 23 habitats had other anopheline larvae and 60 habitats had *Culex* spp (Table 3). This findings suggest that in the *An. funestus* group larvae in their aquatic habitats are more likely to have other different mosquito’s species and organisms similar to the previously studies by Nambunga *et al*., [34] and Dida *et al*., [33].

The association of *An. funestus* group larvae and predators was varying. In particular, this study noted some predators families namely; Notonectidae and Corixidae were bounded in area with low abundance of *An. funestus* group larvae. However the highest number of predators and low number of mosquito larvae could also reflect the direct predation in the aquatic habitats. Whilst Coenagrionidae and Dytiscidae were bounded in the area with higher abundance of *An. funestus* group larvae, showing the positive association between these predators and *An. funestus* group larvae. These could be due to variation in feeding preferences among each predators family. More important, further studies should be done to confirm this because another study has shown that Coenagrionidae are not only significant predators for *Anopheles* larvae but also *Aedes aegypti* larvae [38]. However, the current study did not find a clear and significant association between different predator families like Aeshndae and Belostomatidae with *An. funestus* larvae group.

This study suggests that, for effective malaria vector control, intervention strategies should focus on both, permanent aquatic habitats and temporary/seasonal as well as micro-habitats such as ditches and some man-made wells. The evidence by this study also suggests that these temporary and micro habitats can significantly produce higher numbers of disease transmitting mosquitoes though they limit predator’s colonization abilities. One benefit of utilising biological control is that it may target mosquito species at low densities, it has no impact on non-target organisms and it is simple to use in the field [30].

One of the limitations of this study was that it did not focus on understanding anthropogenic factors and how they might influence the abundance of predators. Though it is very important to assess how both natural and human activities influence abundance of predators, the current study focused on water physicochemical parameters and other physical characteristics of the aquatic habitats. Further studies should also, morphologically identify the aquatic predators to species level using an appropriate identification key. This should help to understand how the aquatic predators are distributed in different aquatic habitats. Although this does not affect our interpretation of results, a cross sectional study doesn’t represent the variation among habitat characteristics over time including the changes in temperature and water physicochemical parameters. A longitudinal study would help capture seasonal variations between predator and prey abundance. Such study may help in the design of novel interventions focussed on this relationship.

## Conclusion

This study demonstrated the existence of common predators in aquatic habitats colonized by *An. funestus* group larvae and factors influencing their abundance. Six predator families were commonly collected; Coenagrionidae, Corixidae, Notonectidae, Amphibians, Aeshndae, and Dytiscidae. The abundance of predator families with *An. funestus* group larvae varied. The only physicochemical parameter influencing *An. funestus* group larvae abundance was dissolved oxygen. Additional studies are needed to demonstrate the efficacy of predators on mosquito larval densities and adult fitness traits. Interventions leveraging the interaction between mosquitoes and predators can be established to disrupt the transmission potential and survival of the *An. funestus* mosquitoes.

## Acknowledgments

We would like to give my great appreciation to community members for their cooperation as well as allowing us to work in their places throughout the data collection period. Also, I would like to extend my sincere gratitude to Outdoor Mosquito Control (OMC) research team members for providing technical and logistical assistance during the study period. We thank Dr. Jacques Derek Charlwood for providing suggestions and reviewing the initial draft of this manuscript.

## List of abbreviations

DO: dissolved oxygen
EC: electrical conductivity
ITNs: Insecticide Treated Nets
IRS: Indoor Residual Spraying
TDS: total dissolved solids
WHO: World Health Organization

## Declarations

### Ethical considerations

Research proposal was presented to the Nelson Mandela Institute of Science and Technology and approval for this study was obtained from the institutional review board of Ifakara Health Institute (Ref: IHI/IRB/No: 13-2022) and from the Medical Research Coordinating Committee (MRCC) at the National Institutes of Medical Research (NIMR) (Ref: NIMR/HQ/R.8a/Vol. IX/3353).

### Consent for publication

The consent of publication this manuscript was obtained from the National Institute for Medical Research (NIMR) Ref. No: NIMR/HQ/P.12 VOL XXXV/61.

## Availability of data and materials

Raw data and material are available from the corresponding author (HHM) upon request

## Author’s contribution

HHM, HSN and EWK conceived the study. HHM, NFK, HSN and EWK organized the field work. HHM, JDK and KSK performed data collection. HHM, LLM and HSN analysed the data. HHM wrote the first draft of the manuscript. JDK, LLM, NFK, HSN and EWK reviewed the manuscript. HSN and EWK reviewed the manuscripts and provided supervision throughout. All authors reviewed and approved the final version of the manuscript.

## References

1. Kileen G, Fillinger U, Bg K. Advantages of larval control for African malaria vectors: low mobility and behavioural responsiveness of immature mosquito stages allow high effective coverage. Malar J. 2002;1:8.

2. Ingabire CM, Hakizimana E, Rulisa A, Kateera F, Van Den Borne B, Muvunyi CM, et al. Community-based biological control of malaria mosquitoes using Bacillus thuringiensis var. israelensis (Bti) in Rwanda: Community awareness, acceptance and participation. Malar J. 2017; 16(1):399.

3. Imbahale SS, Githeko A, Mukabana WR, Takken W. Integrated mosquito larval source management reduces larval numbers in two highland villages in western Kenya. BMC Public Health. 2012;12:1.

4. Fillinger U, Lindsay SW. Larval source management for malaria control in Africa: Myths and reality. Malar. J. 2011; 10:353.

5. Derua YA, Kweka EJ, Kisinza WN, Githeko AK, Mosha FW. Bacterial larvicides used for malaria vector control in sub-Saharan Africa: Review of their effectiveness and operational feasibility. Parasit Vectors. 2019; 12(1):426.

6. Mandal SK, Ghosh A, Bhattacharjee I, Chandra G. Biocontrol efficiency of odonate nymphs against larvae of the mosquito, Culex quinquefasciatus Say, 1823. Acta Trop. 2008; 106(2):109–14.

7. Chandra G, Mandal SK, Ghosh AK, Das D, Banerjee SS, Chakraborty S. Biocontrol of larval mosquitoes by Acilius sulcatus (Coleoptera: Dytiscidae). BMC Infect Dis. 2008;8:1–8.

8. Kweka EJ, Zhou G, Gilbreath TM, Afrane Y, Nyindo M, Githeko AK, et al. Predation efficiency of Anopheles gambiae larvae by aquatic predators in western Kenya highlands. Parasit Vectors. 2011; 4:128.

9. Ohba SY, Kawada H, Dida GO, Juma D, Sonye G, Minakawa N, et al. Predators of Anopheles gambiae sensu lato (Diptera: Culicidae) larvae in Wetlands, Western Kenya: Confirmation by polymerase chain reaction method. J Med Entomol. 2010; 47(5):783–7.

10. Schielke E, Costantini C, Carchini G, Sagnon N, Powell J, Caccone A. Short report: Development of a molecular assay to detect predation on Anopheles gambiae complex larval stages. Am J Trop Med Hyg. 2007; 77(3):464–6.

11. Morales ME, Wesson DM, Sutherland IW, Impoinvil DE, Mbogo CM, Githure JI, et al. Determination of Anopheles gambiae larval DNA in the gut of insectivorous dragonfly (Libellulidae) nymphs by polymerase chain reaction. J Am Mosq Control Assoc. 2003; 19(2):163–5.

12. Aditya G, Saha GK. Predation of the beetle Rhantus sikkimensis (Coleoptera: Dytiscidae) on the larvae of Chironomus Meigen (Diptera: Chironomidae) of the Darjeeling Himalayas of India. Limnologica. 2006;36(4):251–257.

13. Claessen D, Van Oss C, De Roos AM, Persson L. The impact of size-dependent predation on population dynamics and individual life history. Ecology. 2002; 83(6):1660–1675.

14. Eba K, Duchateau L, Olkeba BK, Boets P, Bedada D, Goethals PLM, et al. Bio-control of anopheles mosquito larvae using invertebrate predators to support human health programs in Ethiopia. Int J Environ Res Public Health. 2021; 18(4):1810.

15. Debrah I, Afrane YA, Amoah LE, Ochwedo KO, Mukabana WR, Zhong D, et al. Larval ecology and bionomics of Anopheles funestus in highland and lowland sites in western Kenya. PLoS One. 2021;16(10):e0255321.

16. Ong’Wen F, Onyango PO, Bukhari T. Direct and indirect effects of predation and parasitism on the Anopheles gambiae mosquito. Parasit Vectors. 2020; 13(1):43.

17. Kaindoa EW, Matowo NS, Ngowo HS, Mkandawile G, Mmbando A, Finda M, et al. Interventions that effectively target Anopheles funestus mosquitoes could significantly improve control of persistent malaria transmission in south-eastern Tanzania. PLoS One. 2017;12(5):e0177807.

18. Lwetoijera DW, Harris C, Kiware SS, Dongus S, Devine GJ, McCall PJ, et al. Increasing role of Anopheles funestus and Anopheles arabiensis in malaria transmission in the Kilombero Valley, Tanzania. Malar J. 2014;13:1–10.

19. Mapua SA, Hape EE, Kihonda J, Bwanary H, Kifungo K, Kilalangongono M, et al. Persistently high proportions of plasmodium-infected Anopheles funestus mosquitoes in two villages in the Kilombero valley, South-Eastern Tanzania. Parasite Epidemiol Control. 2022;18:e00264.

20. Coetzee M, Koekemoer LL. Molecular systematics and insecticide resistance in the major African malaria vector anopheles funestus. Annu. Rev. Entomol. 2013; 58:393–412.

21. Charlwood JD. The Ecology of Malaria Vectors. Ecol. Malar. Vectors. 2019.

22. Kaindoa EW, Ngowo HS, Limwagu AJ, Tchouakui M, Hape E, Abbasi S, et al. Swarms of the malaria vector Anopheles funestus in Tanzania. Malar J. 2019; 18(1):29.

23. Ngowo HS, Hape EE, Matthiopoulos J, Ferguson HM, Okumu FO. Fitness characteristics of the malaria vector Anopheles funestus during an attempted laboratory colonization. Malar J. 2021; 20(1):148.

24. Nambunga IH, Ngowo HS, Mapua SA, Hape EE, Msugupakulya BJ, Msaky DS, et al. Aquatic habitats of the malaria vector Anopheles funestus in rural south - eastern Tanzania. Malar J. 2020;19(1):219.

25. Stroud Water Research Center. Identification Guide to Freshwater Macroinvertebrates Major Characteristics of Aquatic Larvae. 2013;6.

26. Gerber A, Gabriel MJM. Aquatic Inverterbrates of South African Rivers: Illustrations. 2002;12.

27. R Development Core Team. R: A language and environment for statistical computing. R Foundation Stat Comput. 2022.

28. Brien JO. Generalized Linear Mixed Models using Template Model Builder. 2022.

29. Arribas R, Touchon JC, Gomez-Mestre I. Predation and competition differentially affect the interactions and trophic niches of a Neotropical amphibian guild. Front Ecol Evol. 2018;6:28.

30. Walker K, Lynch M. Contributions of Anopheles larval control to malaria suppression in tropical Africa: Review of achievements and potential. Med. Vet. Entomol. 2007; (1):2–21.

31. Kamareddine L. The biological control of the malaria vector. Toxins (Basel). 2012;4:748–67.

32. Benelli G, Jeffries CL, Walker T. Biological control of mosquito vectors: Past, present, and future. Insects. 2016;7:1–18.

33. Dida GO, Gelder FB, Anyona DN, Abuom PO, Onyuka JO, Matano AS, et al. Presence and distribution of mosquito larvae predators and factors influencing their abundance along the Mara River, Kenya and Tanzania. Springerplus. 2015; 4:136.

34. Chesson J. Effect of Notonectids (Hemiptera: Notonectidae) on Mosquitoes (Diptera: Culicidae): Predation or Selective Oviposition? Environ Entomol. 1984; 13(2), 531–538.

35. Gilbert JJ, Burns CW. Some observations on the diet of the backswimmer, Anisops wakefieldi (Hemiptera: Notonectidae). Hydrobiologia. 1999;412:111–8.

36. Onen H, Odong R, Chemurot M, Tripet F, Kayondo JK. Predatory and competitive interaction in Anopheles gambiae sensu lato larval breeding habitats in selected villages of central Uganda. Parasit Vectors. 2021; 14(1):420.

37. Banerjee S, Aditya G, Saha N, Saha GK. An assessment of macroinvertebrate assemblages in mosquito larval habitats—space and diversity relationship. Environ Monit Assess. 2010;168:597–611.

38. Collinson NH, Biggs J, Corfield A, Hodson MJ, Walker D, Whitfield M, et al. Temporary and permanent ponds: An assessment of the effects of drying out on the conservation value of aquatic macroinvertebrate communities. Biol Conserv. 1995; 74(2):125–133.

39. Williams P, Whitfield M, Biggs J, Bray S, Fox G, Nicolet P, et al. Comparative biodiversity of rivers, streams, ditches and ponds in an agricultural landscape in Southern England. Biol Conserv. 2004;115(2):329–341.

40. Diabaté A, Dabiré RK, Heidenberger K, Crawford J, Lamp WO, Culler LE, et al. Evidence for divergent selection between the molecular forms of Anopheles gambiae: Role of predation. BMC Evol Biol. 2008;8:5.

41. Carlson J, Keating J, Mbogo CM, Kahindi S, Beier JC. Ecological limitations on aquatic mosquito predator colonization in the urban environment. J Vector Ecol. 2004;29(2):331–9.

42. Mwangangi JM, Mbogo CM, Muturi EJ, Nzovu JG, Githure JI, Yan G, et al. Spatial distribution and habitat characterisation of Anopheles larvae along the Kenyan coast. J Vector Borne Dis. 2007;44(1):44–51.

43. Kenawy MA, Ammar SE, Abdel-Rahman HA. Physico-chemical characteristics of the mosquito breeding water in two urban areas of Cairo Governorate, Egypt. J Entomol Acarol Res. 2013;45:3.

44. Bashar K, Rahman MS, Nodi IJ, Howlader AJ. Species composition and habitat characterization of mosquito (Diptera: Culicidae) larvae in semi-urban areas of Dhaka, Bangladesh. Pathog Glob Health. 2016;110(2):48–6.

45. Clark TM, Vieira MAL, Huegel KL, Flury D, Carper M. Strategies for regulation of hemolymph pH in acidic and alkaline water by the larval mosquito Aedes aegypti (L.) (Diptera; Culicidae). J Exp Biol. 2007; 210:4359–67.

46. Clark TM, Flis BJ, Remold SK. pH tolerances and regulatory abilities of freshwater and euryhaline Aedine mosquito larvae. J Exp Biol. 2004; 207:2297–304.

47. Akeju AV, Olusi TA, Simon-Oke IA. Effect of physicochemical parameters on Anopheles mosquitoes larval composition in Akure North Local Government area of Ondo State, Nigeria. J Basic Appl Zool. 2022;83:34.

48. Obi OA, Nock IH, Adebote DA. Ecology of Preimaginal Culicine Mosquitoes in Rock Pools on Inselbergs Within Kaduna State, Nigeria. J Mosq Res. 2019; 9(5):35–48.

49. Chaiphongpachara T, Yusuk P, Laojun S, Kunphichayadecha C. Environmental Factors Associated with Mosquito Vector Larvae in a Malaria-Endemic Area in Ratchaburi Province, Thailand. Sci World J. 2018; 2018:4519094.

50. Spartial Variation in Physicochemical Characteristics of Wetland Rice Fields Mosquito Larval Habitats in Minna, North Central Nigeria. AEMS. 2015;10:53–56.

51. Johansson F, Brodin T. Effects of fish predators and abiotic factors on dragonfly community structure. J Freshw Ecol. 2003;18(3):415–423.

52. Gillooly JF, Brown JH, West GB, Savage VM, Charnov EL. Effects of size and temperature on metabolic rate. Science. 2001;293(5538):2248–2251.

53. Teoh ML, Chu WL, Phang SM. Effect of temperature change on physiology and biochemistry of algae: A review. Malaysian J. Sci. 2010;29(2):82–97.

54. Getachew D, Balkew M, Tekie H. Anopheles larval species composition and characterization of breeding habitats in two localities in the Ghibe River Basin, southwestern Ethiopia. Malar J. 2020;19:1–13.

55. Musonda M, Sichilima A. The effect of total dissolved solids, salinity and electrical conductivity parameters of water on abundance of anopheles mosquito larvae in different breeding sites of kapiri mposhi district of Zambia. Int J Sci Technol Res. 2019;8: 2277–8616.

56. Amini M, Hanafi-Bojd AA, Aghapour AA, Chavshin AR. Larval habitats and species diversity of mosquitoes (Diptera: Culicidae) in West Azerbaijan Province, Northwestern Iran. BMC Ecol. 2020;20(1):60.

57. Oyewole IO, Momoh OO, Anyasor GN, Ogunnowo AA, Ibidapo CA, Oduola OA, et al. Physico-chemical characteristics of Anopheles breeding sites: Impact on fecundity and progeny development. African J Environ Sci Technol. 2009;3:447–52.

58. Abai MR, Saghafipour A, Ladonni H, Jesri N, Omidi S, Azari-Hamidian S. Physicochemical characteristics of larval habitat waters of mosquitoes (Diptera: Culicidae) in qom province, central iran. J Arthropod Borne Dis. 2016;10(1):65–77.

